# Exploratory growth in *Streptomyces venezuelae* involves a unique transcriptional program, enhanced oxidative stress response, and profound acceleration in response to glycerol

**DOI:** 10.1101/2021.12.22.473798

**Authors:** Evan M. F. Shepherdson, Tina Netzker, Yordan Stoyanov, Marie A. Elliot

## Abstract

Exploration is a recently discovered mode of growth and behaviour exhibited by some *Streptomyces* species that is distinct from their classical sporulating life cycle. While much has been uncovered regarding initiating environmental conditions and the phenotypic outcomes of exploratory growth, how this process is coordinated at a genetic level remains unclear. We used RNA-sequencing to survey global changes in the transcriptional profile of exploring cultures over time in the model organism *Streptomyces venezuelae*. Transcriptomic analyses revealed widespread changes in gene expression impacting diverse cellular functions. Investigations into differentially expressed regulatory elements revealed specific groups of regulatory factors to be impacted, including the expression of several extracytoplasmic function (ECF) sigma factors, second messenger signalling pathways, and members of the *whiB*-like (*wbl*) family of transcription factors. Dramatic changes were observed among primary metabolic pathways, especially among respiration-associated genes and the oxidative stress response; enzyme assays confirmed that exploring cultures exhibit an enhanced oxidative stress response compared with classically growing cultures. Changes in expression of the glycerol catabolic genes in *S. venezuelae* led to the discovery that glycerol supplementation of the growth medium promotes a dramatic acceleration of exploration. This effect appears to be unique to glycerol as an alternative carbon source and this response is broadly conserved across other exploration-competent species.

## INTRODUCTION

The soil is a highly heterogeneous environment. Essential growth factors like oxygen, carbon, and iron can be present in widely varying levels (1–4), and these nutrient levels can be shaped by biotic (e.g. microbial population density) and abiotic factors (e.g. water saturation), which are themselves dynamic. Bacterial survival in this complex, fluctuating environment requires both genetic and metabolic flexibility, and an ability to effectively compete or cooperate with neighbouring microbes.

In the soil, *Streptomyces* are a genus of ubiquitous Gram-positive bacteria that epitomize both metabolic and developmental adaptability. As saprophytic organisms, *Streptomyces* harbour a vast repertoire of catabolic enzymes that allow them to make use of a diverse set of carbon sources, including monomeric carbon sources such as glycerol, mannitol, arabinose, and galactose, through to complex polysaccharides like chitin and cellulose (5). Similarly, the genomes of different *Streptomyces* species have revealed remarkable metabolic capabilities, in the form of abundant biosynthetic gene clusters that direct the production of diverse specialized metabolites, often with little overlap between species (6). These specialized metabolites are often small molecules with potent biological activity, and include compounds with the ability to inhibit the growth of bacteria and fungi (e.g. antibiotics) (7, 8), promote iron acquisition (e.g. siderophores) (9), and influence community behaviours (e.g. quorum sensing molecules) (10, 11). This rich library of chemical effectors is proposed to equip the streptomycetes with a powerful repertoire of tools that facilitate either competitive or cooperative interactions with other microbes.

Complementing the robust metabolic capabilities of the streptomycetes is a remarkable developmental flexibility. The classical *Streptomyces* life cycle begins with spore germination, and proceeds through the growth of branching filamentous hyphae, establishing a metabolically active vegetative mycelium. Nutrient depletion or other stressors induce a coordinated switch into reproductive growth, which initiates with the raising of aerial hyphae (12). Further maturation events promote the differentiation of these aerial structures into chains of metabolically dormant spores. These spores are resistant to many abiotic stresses, and can be released from their chains and dispersed to new environments (13).

Recently, it has been shown that some *Streptomyces* species can abandon their canonical, classical life cycle when grown on solid surfaces, and enter an alternative growth mode called ‘exploration’. Exploratory growth is phenotypically distinct from classical growth: colonies expand rapidly outward as a vegetative-like mycelium and develop a dense network of wrinkles at the centre of the biomass (14). In *Streptomyces venezuelae*, exploration is initiated in response to specific environmental cues. Stimulation of exploration was originally identified as a competitive response by *S. venezuelae* to co-culture with the yeast *Saccharomyces cerevisiae* on a rich, glucose-containing medium (yeast extract – peptone – dextrose; YPD) (14). An analogous response can be initiated in the absence of yeast when the same rich medium lacks glucose (yeast extract – peptone; YP), highlighting a strong repressive effect of glucose on exploration (14).

Concurrent with morphological development, exploring colonies reshape the physicochemical properties of their surrounding environment through the emission of trimethylamine, a small basic volatile compound (14). The airborne diffusion of trimethylamine away from the colony promotes a steady rise in local pH, which in turn confers a competitive advantage to exploring *Streptomyces* species. It promotes the exploration of nearby *S. venezuelae* cultures growing on rich, glucose-containing medium (typically an exploration-repressive condition), and simultaneously lowers the bioavailability of iron by forming poorly soluble hydroxides (15). This self-imposed iron limitation both strengthens the response of exploring *S. venezuelae* cultures and inhibits the growth of other microbes in the vicinity.

While the environmental triggers of exploration initiation and propagation are becoming clearer, there is much that remains to be understood about this unusual microbial behaviour and the factors that influence it. Exploration in *Streptomyces* appears to be largely independent of regulators that control classical development (14). This raises an interesting possibility that exploration is encoded as an independent genetic program, with both defined regulatory circuits and distinct metabolic pathways that may be activated or down-regulated over time. To better understand the pathways that drive *Streptomyces* exploration, we examined gene expression changes over time in *S. venezuelae* exploring cultures using RNA-sequencing. We observed a global reprogramming of transcription during the maturation of an exploring colony, with notable upregulation of genes associated with primary metabolism, respiration, and the oxidative stress response. A number of regulatory elements belonging to different functional categories also exhibited changes in expression over time, including known regulators of classical development, extracytoplasmic function (ECF) sigma factors, and second messenger systems. We further discovered that glycerol supplementation of the growth medium radically enhanced the exploration response in *S. venezuelae* and altered the metabolic output of these cells. This phenomenon was unique to glycerol, as a similar response could not be induced by supplementation with other alternative carbon sources. Conversely, the response to glycerol was not unique to *S. venezuelae*, as multiple exploration-competent wild *Streptomyces* isolates exhibited analogous enhanced exploratory growth in the presence of glycerol.

## RESULTS

### Gene expression is globally reprogrammed during exploration on YP medium

We previously observed that *S. venezuelae* exploration occurs independently of many of the key regulators of classical development (i.e. the *bld* and *whi* genes), suggesting that a different suite of signalling pathways may function to coordinate the switch into exploration. To gain insight into the genes that are activated and repressed during *S. venezuelae* exploration, we performed RNA-sequencing on transcripts isolated from *S. venezuelae* colonies growing on exploration-promoting medium (YP) over time. Representative timepoints were selected based on colony morphology to capture each phase of the exploration process: early (colony establishment), mid (establishment of a ‘core’ region and start of exploration), and late (robust exploration) (**Figure 1A**). Analysis of gene expression data between the early and late timepoints revealed over 1,000 genes that were differentially expressed and were distributed across the chromosome (**Figure 1B**). Reducing the complexity within the data to Clusters of Orthologous Groups (COGs), we saw a high frequency of differential expression for genes participating in primary metabolism (COG terms: carbohydrate transport and metabolism; amino acid transport and metabolism; and inorganic ion transport and metabolism) (**Figure 1C**). When identifying target genes for follow-up validation studies, we first focused on operons that showed consistent trends in expression over time and genes that participated in similar pathways.

**Figure 1.**
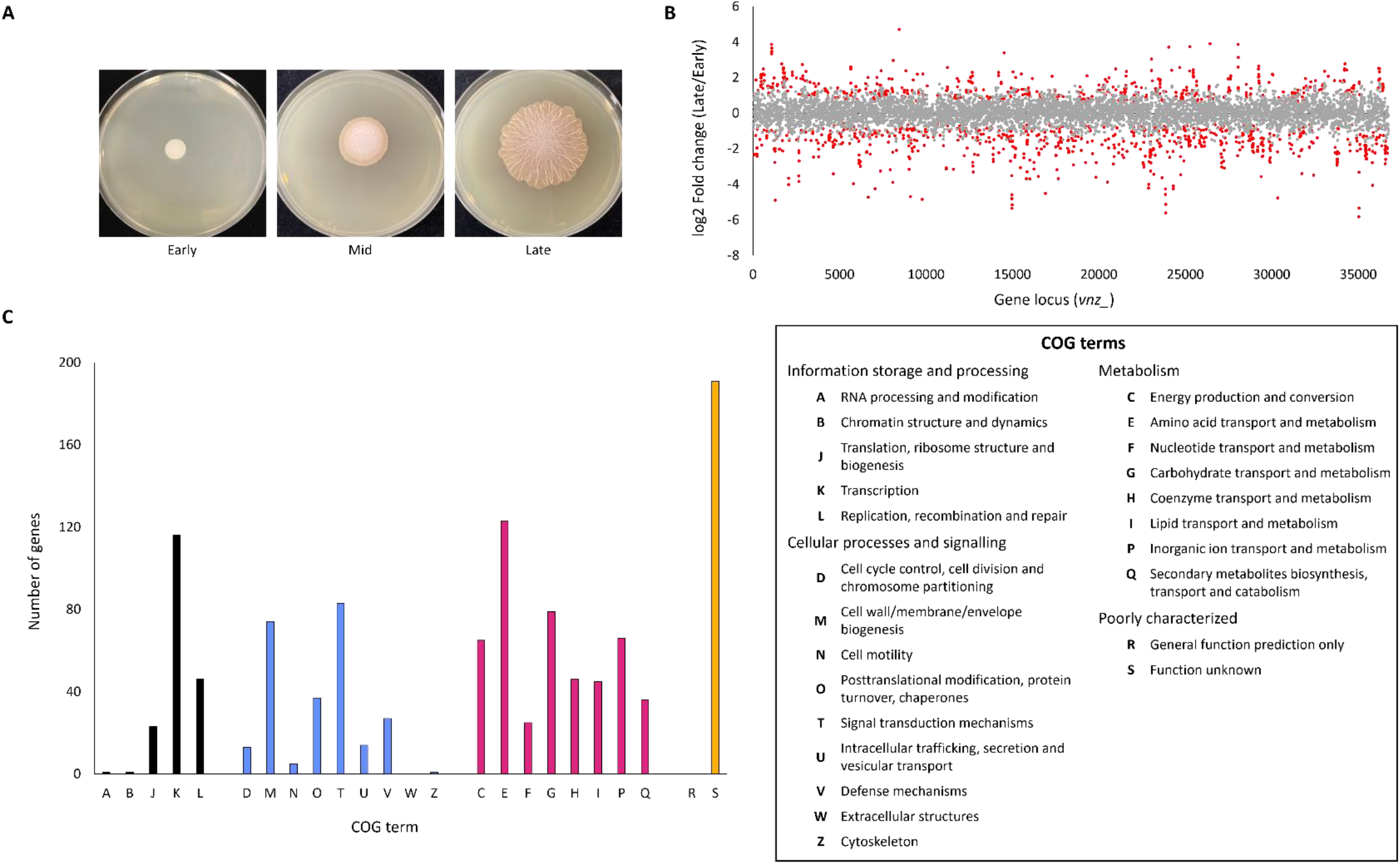
RNA-sequencing of cultures exploring on YP over time. **A)** Representative colonies chosen as timepoints for RNA isolation and sequencing. Early, mid, and late cultures were grown for 2, 5, and 9 days, respectively. **B)** Differential gene expression over time in YP exploring cultures. The fold change in expression of each gene between the early and late timepoint was plotted, and these were arranged by their position on the *S. venezuelae* chromosome. Data points that are statistically significant are shown in red (p < 0.01). **C)** Genes that showed statistically significant (p < 0.01) differential expression between early and late timepoints were assigned Cluster of Orthologous Groups (COG) identifiers and their distributions were plotted. A legend for the corresponding COG terms is presented to the right of the graph.

Among the most statistically significant differentially expressed genes were those involved in inorganic nitrogen (highest transcript levels early) and sulfur metabolism (highest transcript levels late) (**Figure 2A**). Therefore, we tested the exploration response when concentrations of these nutrients were altered in exploration-promoting conditions. We supplemented YP with nitrate, sulfite, and sulfate at a range of concentrations (10 μM-1 mM), and found that in all instances, the effect on *S. venezuelae* exploration was negligible (**Figure 2B**). From this we concluded that nitrogen and sulfur acquisition were likely important, but that sufficient concentrations of these nutrients could be obtained from the growth medium such that additional supplementation had no effect.

**Figure 2.**
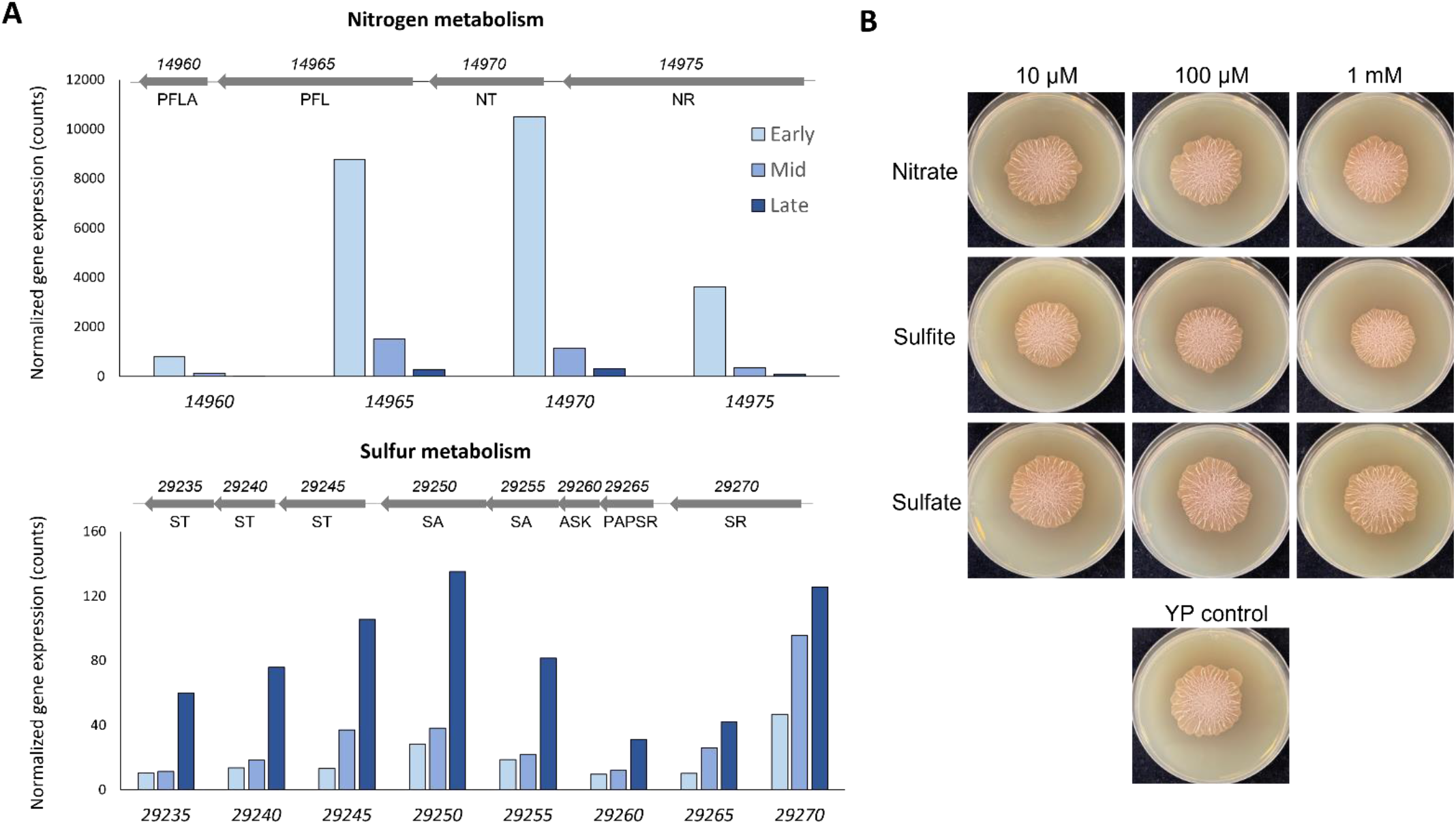
Effects of inorganic nitrogen and sulfur during exploration on YP. **A)** Gene expression levels in YP over time for two regions of the *S. venezuelae* chromosome with predicted function in inorganic nitrogen (*vnz_14960-14975*, top) and sulfur (*vnz_29235-29270*, bottom) metabolism. Gene organization for each region is depicted above each graph, with *vnz* gene identification numbers indicated above and gene annotations below each arrow. *PFLA* pyruvate-formate lyase activating enzyme; *PFL* pyruvate-formate lyase; *NT* nitrate/nitrite transporter; *NR* nitrite reductase; *ST* sulfate transporter; *SA* sulfate adenylyltransferase; *ASK* adenylyl-sulfate kinase; *PAPSR* phosphoadenosine phosphosulfate reductase; *SR* sulfite reductase. **B)** Effects of nitrogen and sulfur supplementation on exploration. *S. venezuelae* was spotted to YP medium supplemented with varying concentrations of nitrate, sulfite, and sulfate and imaged after 7 days of growth.

### Exploration on YP is associated with a strong respiration and oxidative stress response

Analysis of our RNA-sequencing data further suggested that energy production and respiration pathways were similarly induced over the course of exploration. We saw strong transcriptional upregulation of genes encoding ATP synthase subunits, alongside genes encoding iron-sulfur cluster assembly proteins, which produce essential cofactors responsible for conducting electrons through the supercomplexes of the electron transport chain (**Figure 3A-B**). Accordingly, as the respiratory chain is a noted source of intracellular reactive oxygen species (16), we saw a corresponding transcriptional activation of the genes encoding the oxidative stress-responsive catalases and superoxide dismutases (**Figure 3C**).

**Figure 3.**
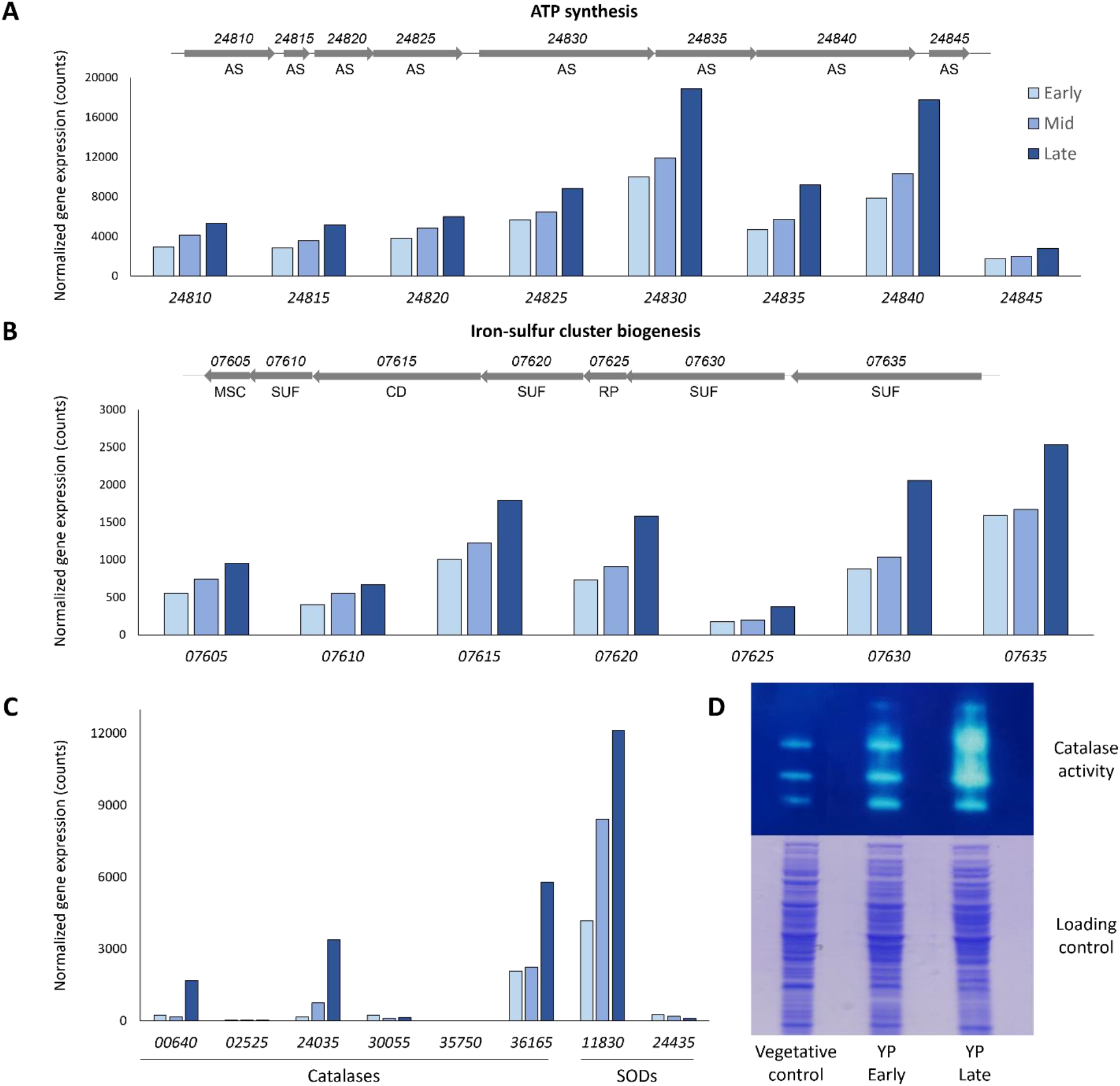
Upregulation of respiration-associated and oxidative stress response genes during exploration on YP. **A)** Expression over time for genes participating in ATP synthesis (*vnz_24810-24845*). **B)** Expression over time for genes participating in iron-sulfur cluster biogenesis (*vnz_07605-07635*). **(A, B)** Gene organization for each region is depicted above each graph, *vnz* gene identification numbers are given above and gene annotations are given below each arrow. *AS* ATP synthase subunit; *MSC* metal-sulfur cluster biosynthetic enzyme; *SUF* sulfur assimilation (SUF) system iron-sulfur cluster assembly protein; *CD* cysteine desulfurase; *RP* Rieske (2Fe-2S) protein. **C)** Gene expression levels in YP-grown cultures over time for predicted catalase and superoxide dismutase (SOD) genes encoded by *S. venezuelae*. **D)** Catalase enzyme activity was assayed by native PAGE (top) using proteins from cultures harvested at early and late timepoints from exploring cells grown on YP, together with extracts from classical vegetatively-grown cultures. Coomassie-stained SDS-PAGs (bottom) of the same samples served as a loading control for total protein content.

To determine whether increased transcription of these oxidative stress-protective enzyme-encoding genes was correlated with increased enzyme activity, we assessed catalase activity in exploring cultures over time. Native PAGE was used to resolve distinct catalase species from whole cell lysates harvested from early- and late-stage YP exploring cultures, alongside a vegetative (non-exploring) growth control. We found that at our early timepoint (before exploration had initiated), activity levels were roughly equivalent to those from classical, vegetative growth conditions (**Figure 3D**). However, at the late timepoint (during robust exploratory growth) there was a pronounced increase in catalase activity, suggesting that these cells had an enhanced capacity to manage reactive oxygen species, compared with early and vegetatively-grown cultures.

To assess whether increased catalase activity was a pre-requisite for robust exploration, we generated strains of *S. venezuelae* in which the most highly-expressed catalase gene in our transcriptional dataset (*vnz_36165*) was either deleted or overexpressed. When these strains were grown in parallel with their equivalent wild type control strain on YP, there were subtle differences in colony architecture compared with the wild type; however, the overall rate of exploration was not significantly impacted for either the deletion or overexpression strain (**Supplemental Figure 1**).

**Supplemental Figure 1.**
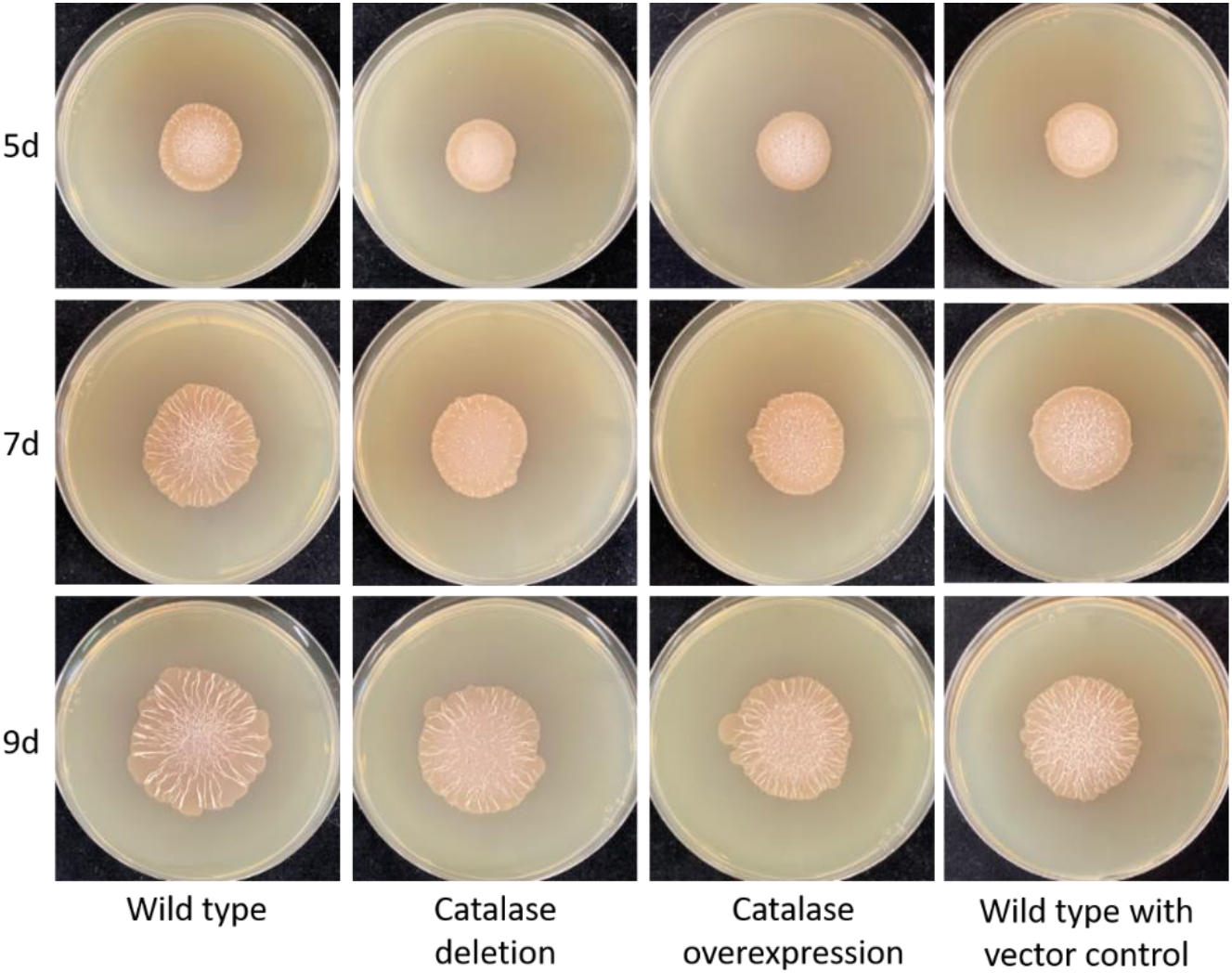
Modulating catalase levels during exploration on YP. Strains in which the catalase-encoding *vnz_36165* was deleted or overexpressed were spotted alongside wild type *S. venezuelae* alone (left) or carrying an empty vector control for the overexpression construct (right), on YP medium. Images were taken of representative colonies at three timepoints.

### Multiple mechanisms of transcriptional control are differentially expressed during exploration

To identify regulatory elements that may function in coordinating the exploration response, we extracted differentially expressed genes from the COG analysis (**Figure 1C**) that had been assigned to the terms ‘transcription’ or ‘signal transduction mechanisms’ (**Table S1**). As the RNA-sequencing data had suggested that exploratory growth was concomitant with a broad reprogramming of gene expression, we initially prioritized regulators that we expected to function globally in the cell rather than modulating pathway-specific processes. To this end, we identified multiple functional categories of regulators that could fulfill this role, including well-characterized transcription factors that participate in classical development, extracytoplasmic function (ECF) sigma factors, and second messenger-responsive proteins (**Table 1; Table S1** – provided as a supplementary data file).

**Table 1.** Global regulators that are differentially expressed over the course of exploration.

| Gene/Product | Log<sub>2</sub>Fold Change (Early/Late) | p-value |
| --- | --- | --- |
| **Regulators of classical development** |  |  |
| *bldM* (vnz_22005) | 1.12 | $2.76 \times 10^{-3}$ |
| *bldN* (vnz_15655) | 2.50 | $1.01 \times 10^{-4}$ |
| *whiH* (vnz_27205) | 2.70 | $1.03 \times 10^{-11}$ |
| **WhiB-like proteins** |  |  |
| *whiB* (vnz_13645) | 1.37 | $1.29 \times 10^{-6}$ |
| *wblA* (vnz_16495) | 3.56 | $2.35 \times 10^{-36}$ |
| *wblE* (vnz_24255) | 1.20 | $1.68 \times 10^{-4}$ |
| *vnz_00590* | -2.49 | $1.15 \times 10^{-4}$ |
| **ECF sigma factors** |  |  |
| *sigE* (vnz_15840) | 3.30 | $7.46 \times 10^{-37}$ |
| *sigQ* (vnz_22610) | 3.02 | $1.82 \times 10^{-13}$ |
| **Second messenger-responsive proteins** |  |  |
| *eshA* (vnz_35035) | 5.82 | $1.14 \times 10^{-33}$ |

As the majority of these regulators were more highly expressed early in the exploration process, we wondered whether any of these systems were important for relaying signals needed to initiate exploration within the cell. In the set of regulators that overlapped with those coordinating classical development, we identified *bldM, bldN*, and *whiH*. BldN is an alternative sigma factor that directs the expression of multiple development-specific genes including *bldM*, which encodes a transcription factor that functions both alone and in conjunction with WhiI to control the transcription of genes needed for reproductive growth (17, 18). We know from previous work that mutants defective in *bldN*, and to a lesser extent *bldM*, are capable of exploring, but do so at a reduced rate (14). WhiH acts at a later stage in classical development and is important for spore formation (12). Deletion of its associated gene has previously been shown to have a negligible impact on the progression of exploration (14).

The WhiB-like (Wbl) proteins comprise a subcategory of classical development regulators. This family of typically short proteins (< 150 amino acids) is specific to the *Actinobacteria*, and their associated genes are often found in multiple copies (e.g. 11 paralogs are encoded within the *S. venezuelae* chromosome). While the function of many of these proteins remain unknown, three members (WhiB, WhiD, and WblA) have been shown in different *Streptomyces* species to act as transcription factors that coordinate distinct stages of reproductive growth (19). A defining feature shared by Wbl proteins is the conserved presence of a [4Fe-4S] cluster that is necessary for protein function and is sensitive to decomposition by both O_2_ and NO (19). Although we noted strong upregulation of protein-coding genes functioning in iron-sulfur cluster biosynthesis during exploration (**Figure 3**), deleting the founding member of the Wbl family, *whiB*, still allowed for robust exploration. Notably, known targets of WhiB (including *ftsZ* and *ftsW*) were not found to be differentially expressed over the course of exploration.

The third set of global regulators we identified in our differential expression analyses were the extracytoplasmic sigma factors *sigE* and *sigQ*. As their name implies, these sigma factors are commonly activated upon sensing an environmental stimulus. Often, ECF sigma factors are regulated post-translationally by anti-sigma factors. Curiously, SigE and SigQ are two exceptions; instead, they are under the transcriptional control of genome-adjacent two-component systems *cse(A)BC* and *afsQ1/Q2*, respectively (20–23). The SigE regulon has recently been defined in *Streptomyces coelicolor* by chromatin immunoprecipitation; many of its target genes contribute to cell envelope function and integrity (24). Much less is known about SigQ; it is broadly conserved among *Streptomyces* species and mutational studies in *S. coelicolor* have implicated it in both development and antibiotic production (22, 25). Given that less was known about SigQ, and given its conserved nature, we were interested in probing its role in exploration. We created both *sigQ* deletion and overexpression strains, and compared their growth to a wild type strain on YP medium. Robust exploration was observed for all strains (**Figure 4A**). Subtle differences were noted for the *sigQ* overexpression strain relative to wild type, including a reduced rate of expansion, coupled with more wrinkling and apparent biomass formation in the core region of the colony.

**Figure 4.**
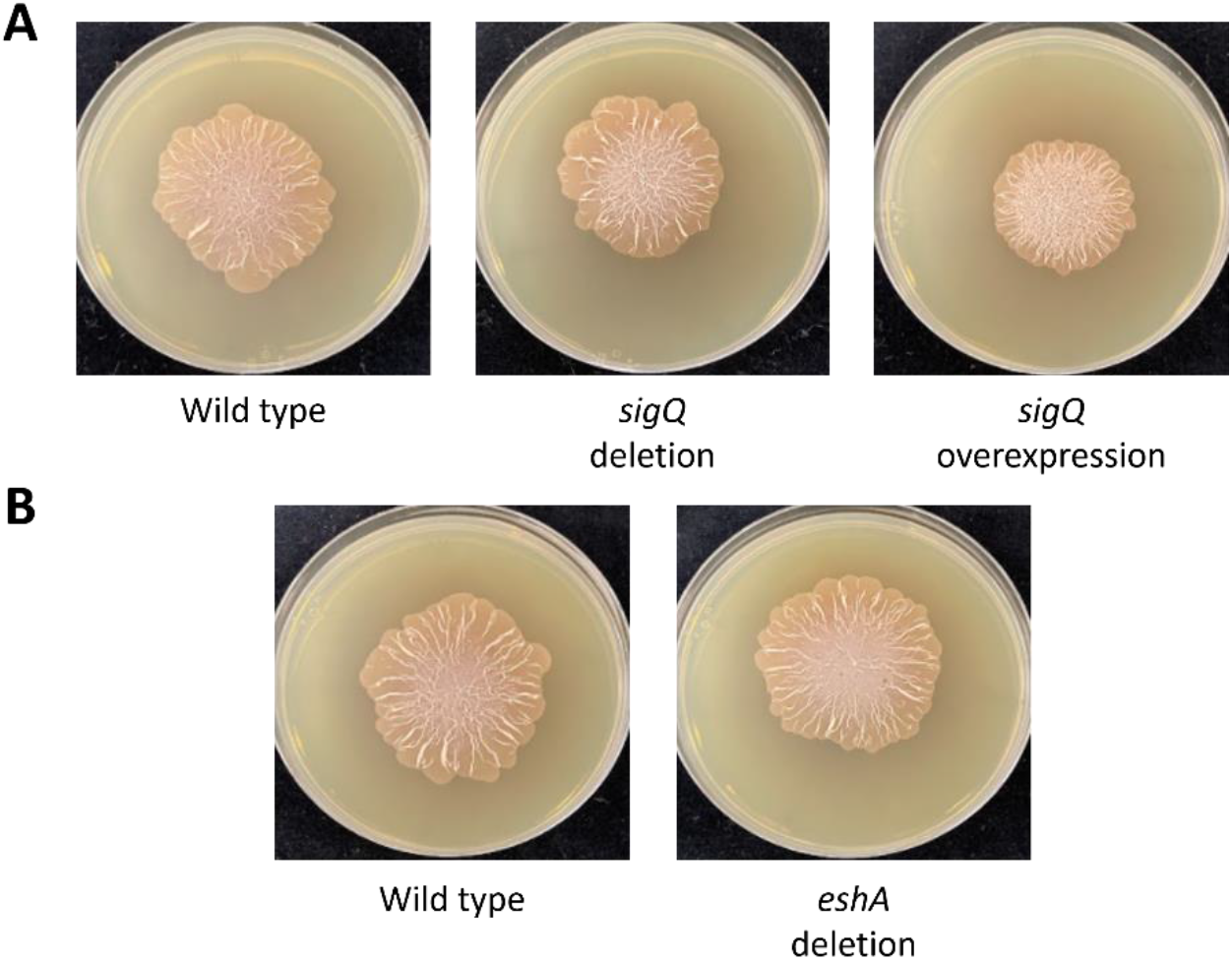
Contribution of differentially expressed global regulators to exploration. Mutant strains of the global regulators *sigQ* (**A**) and *eshA* (**B**) were spotted to YP medium, alongside wild type *S. venezuelae*, and were assessed for exploratory growth. Images were taken after 9d of growth.

The final category of regulators identified as being differentially expressed was the second messenger-binding proteins. EshA is a conserved cAMP-binding protein that has been implicated in different processes depending on the species. In *S. coelicolor, eshA* deletion results in reduced production of the antibiotic actinorhodin, while development is unaffected (26). Conversely, in *S. griseus, eshA* deletion abolishes aerial mycelium formation, and also reduces the production of the antibiotic streptomycin (27). As *eshA* exhibited one of the largest fold changes in expression over time during exploration and has previously been tied to developmental transitions, we wondered if EshA may also have a regulatory role in exploring cultures. We found that an *eshA* mutant was indistinguishable from the wild type with respect to its exploration capabilities (**Figure 4B**).

### Glycerol profoundly affects exploration

We had previously established that carbon source (glucose) had a strong effect on exploration. When expanding our analysis of transcriptional regulators to include those that had more pathway-specific functions, we identified significant differential expression for the regulator of glycerol catabolism, *gylR*, with similar observed trends for its downstream regulon members. The *gyl* region consisted of an operon of three genes (*gylFKD*) that encoded a permease, a glycerol kinase, and a glycerol-3-phosphate dehydrogenase, respectively, (**Figure 5A**); expression of *gylR* and the *gylFKD* operon was highest at the earliest timepoint, and expression rapidly tapered off as exploration proceeded (**Figure 5B**). We wondered how cells would respond if their growth medium was supplemented with glycerol, and if this might delay the onset of exploration. To test this, we compared the growth of wild type *S. venezuelae* on YP and YP supplemented with glycerol (YPG). Unexpectedly, the addition of glycerol dramatically accelerated *S. venezuelae* exploration (**Figure 5C**). Compared to their YP-grown counterparts, YPG explorers grew significantly more rapidly (**Figure 5D; Supplemental Video 1**); colonies initially spotted at the centre of a standard 10 cm Petri plate had expanded to cover the entire plate within 7-9 days of growth; such a colony size was not typically observed for YP explorers even with extended incubation times. YPG-grown colonies also developed a distinct colony morphology compared with YP-grown cultures, having a much more intricate network of wrinkles. Growth on YPG further yielded significantly more condensation within the plates compared with YP-grown cultures, suggesting enhanced respiration rates (**Supplemental Video 1**). Beyond the differences in exploration growth rate and colony architecture, we also found that YPG-grown colonies began secreting a bright orange pigment into the underlying medium after approximately 5 days of growth (**Figure 5E**).

**Figure 5.**
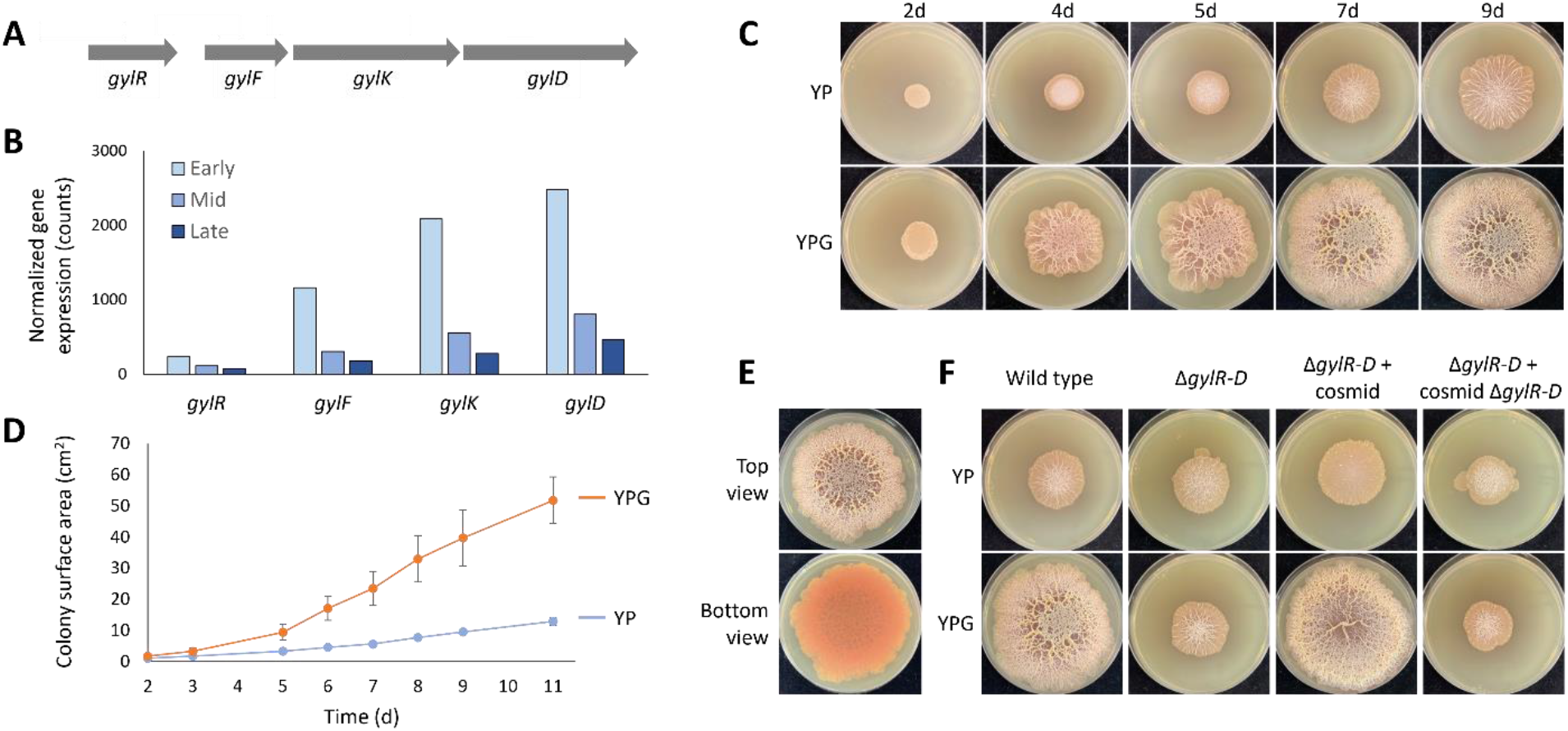
Effect of glycerol on exploration. **A)** Organization of the glycerol-utilization operon in *S. venezuelae*. The three genes of the *gylFKD* operon are transcribed from a single promoter. **B)** Transcript levels in YP over time for the glycerol catabolic gene cluster. **C)** Images of representative *S. venezuelae* colonies spotted on YP and YPG at 2-9 days after inoculation. **D)** Solid media growth curves comparing the rate of surface area expansion for *S. venezuelae* grown on YP and YPG. The average of six replicates was plotted for each condition. Error bars represent one standard deviation. **E)** Representative images of a YPG-grown colony from the top and bottom side of the plate show secretion of an orange pigment to the underlying agar. Images were taken after 7d of growth. **F)** Growth of wild type *S. venezuelae*, a glycerol operon deletion mutant (Δ*gylR-D*), a complemented strain (Δ*gylR-D* + cosmid), and a complementation vector control strain (Δ*gylR-D* + cosmid Δ*gylR-D*) were spotted to YP and YPG. Representative images were taken after 7d of growth.

To confirm that these changes in *S. venezuelae* exploration were mediated by glycerol uptake and metabolism, we generated a mutant strain in which *gylR* and the *gylFKD* operon were replaced with an apramycin resistance cassette. When inoculated on YPG plates, the mutant grew as if it were “blind” to glycerol, phenotypically resembling the wild type strain grown on YP (**Figure 5F**). The dramatic YPG exploration phenotype could be restored to the mutant by re-introducing a wild type copy of these genes *in trans* (**Figure 5F**).

Given the remarkable changes in exploration that stemmed from growth on glycerol, we were curious if the effects were specific to glycerol or if a similar effect could be conferred by supplementing with any alternative carbon source. To test this, we grew wild type *S. venezuelae* on YP (no glucose-containing) plates, supplemented with a variety of carbon sources (**Supplemental Figure 2**). Of the eight carbon sources tested, none enhanced exploration in the way that glycerol did.

**Supplemental Figure 2.**
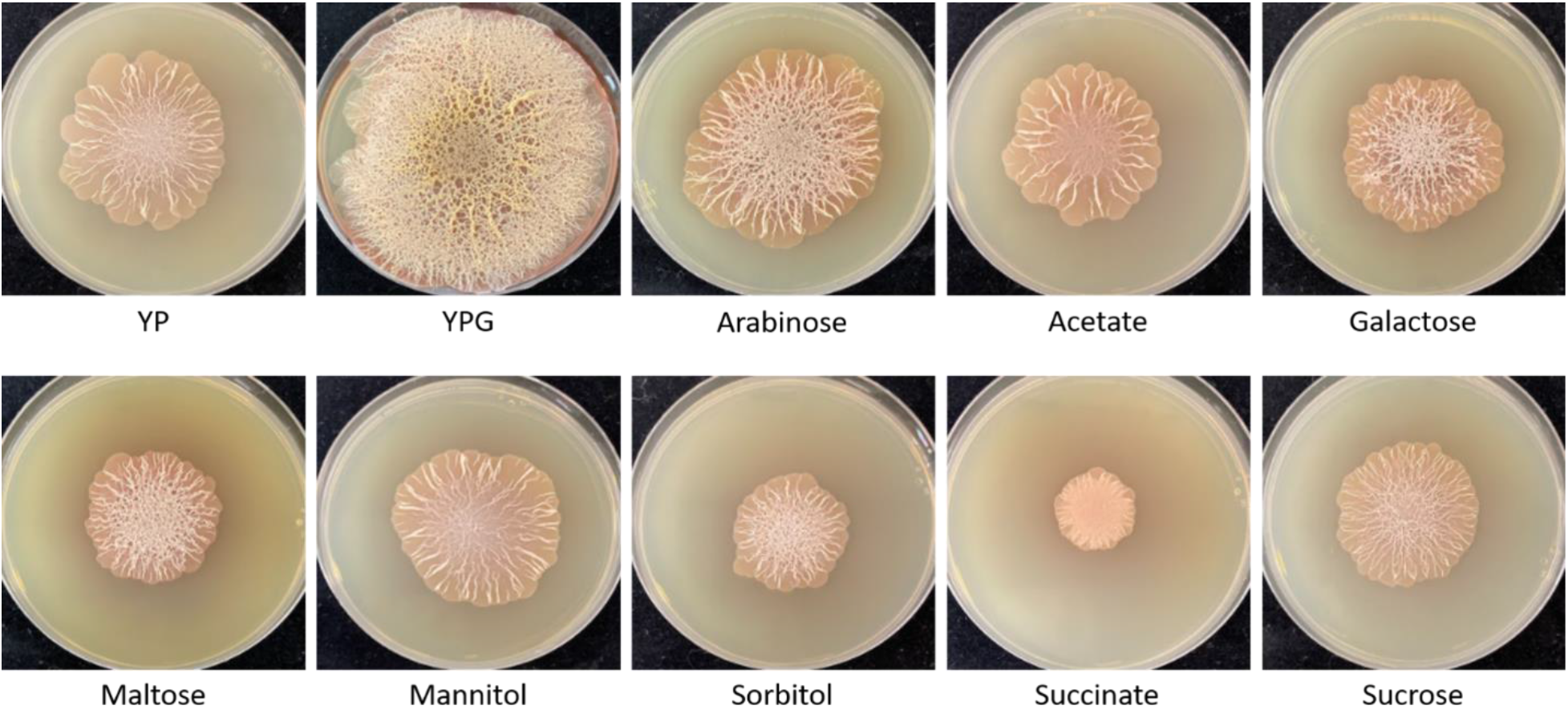
Effects of alternative carbon sources on exploration. *S. venezuelae* was spotted to YP supplemented with the indicated carbon sources at a final concentration of 2% (w/v). Images of representative colonies were taken after 9d of growth.

Similarly, we were interested to know whether the glycerol effects on exploration were specific to *S. venezuelae* or if this effect was also observed for other streptomycetes. As the exploration response is not universally conserved among streptomycetes (or may be triggered by as yet undiscovered conditions for some species), we assembled a panel of 21 wild *Streptomyces* isolates that showed robust exploration on YP. For these strains, glycerol supplementation consistently resulted in colonies with increased wrinkling (21/21) and pigmentation (17/21) (**Figure 6**). Glycerol supplementation was also frequently associated with accelerated exploration; over 60% (13/21) of these strains yielded colonies that were larger on YPG relative to YP (*e.g*. WAC 5485 – **Figure 6**). A full summary of phenotypes for the tested strains can be found in **Table S2** (provided as a supplementary data file). In all, these data suggest that glycerol can dramatically alter exploration dynamics, colony architecture, and metabolism in diverse streptomycetes.

**Figure 6.**
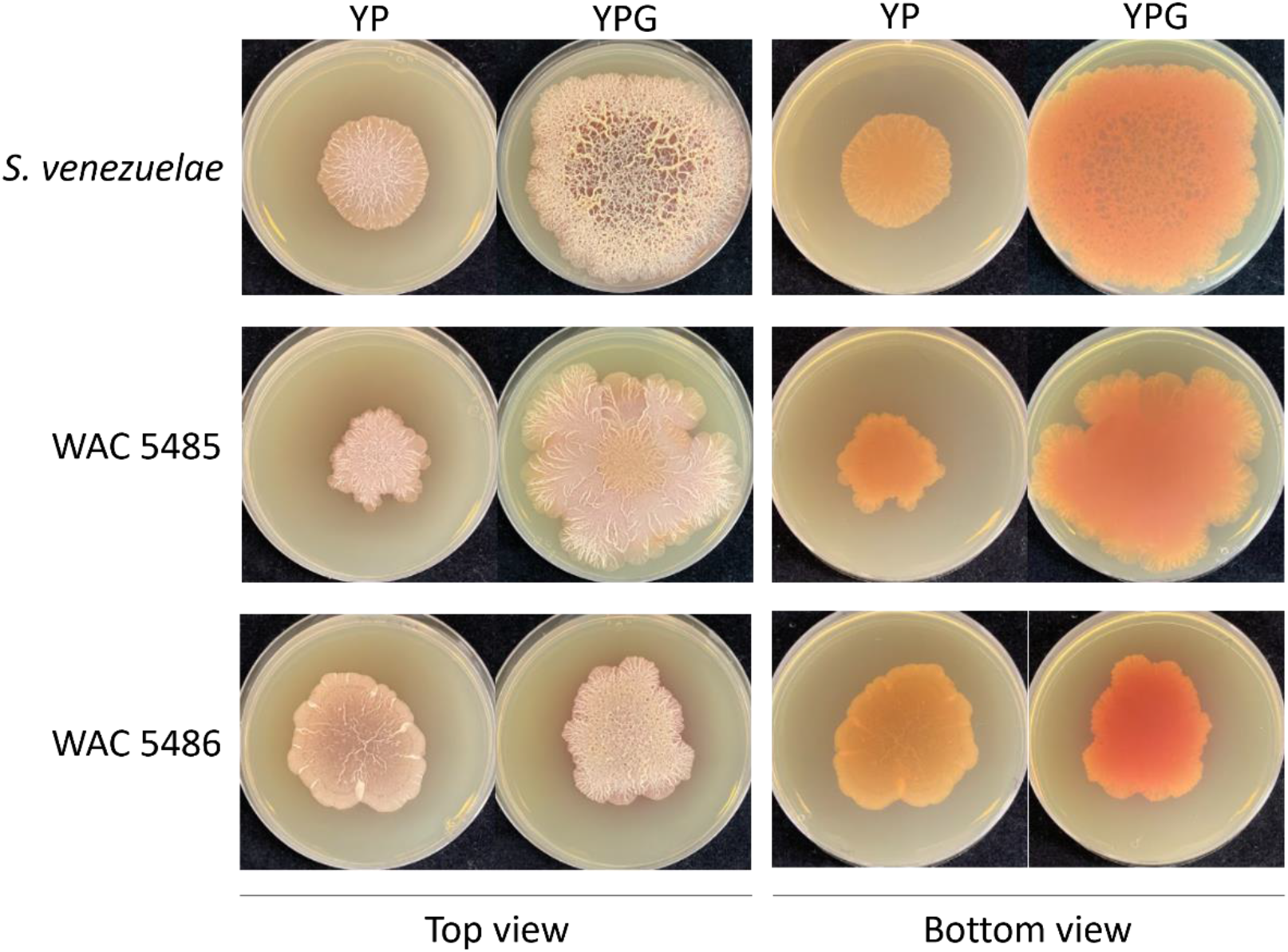
Effects of glycerol on exploration in other *Streptomyces* species. Different wild *Streptomyces* isolates from the Wright Actinomycete Collection (WAC) were spotted alongside *S. venezuelae* on YP and YPG media. Representative colonies of select strains were photographed following 7d of growth.

## DISCUSSION

Our investigation into the transcriptional profile of exploring *S. venezuelae* revealed that there was a global remodeling of gene expression over time in YP-grown cells. Just as changes were not localized to specific regions of the chromosome, the functions of differentially expressed genes were similarly diverse. We saw notable up- and down-regulation of genes participating in organic and inorganic nutrient acquisition, primary metabolism, and cellular defense, alongside other cellular functions and pathways. Known regulators that participate in classical development had at most, subtle effects on exploratory growth, implying that additional strategies must contribute to the control of this behaviour. While we were unable to implicate a specific regulatory element in coordinating exploration, there were many regulatory genes whose expression changed significantly over the course of an exploration growth cycle. It will be interesting to test the effects of some of the less well-characterized regulators on exploration, both alone, and in combination with others.

We found exploration was associated with an enhanced oxidative stress response, involving the upregulation of multiple superoxide dismutase- and catalase-encoding genes, and significantly increased catalase enzyme activity. Manipulating the levels of the most highly expressed catalase had minimal effects on exploration, suggesting considerable functional redundancy shared amongst these related enzymes. Notably, one of the regulatory genes whose expression decreased as exploration proceeded was *wblA*. The WblA-class of regulators has been shown to negatively regulate the oxidative stress response in *S. coelicolor* (51), *Corynebacterium glutamicum* (52), and *Mycobacterium tuberculosis* (53). It is conceivable that WblA has a similar role in *S. venezuelae*, and its downregulation during exploration is needed to mount an effective oxidative stress response.

Our work here, alongside previous investigations into exploration (14, 15), suggests that increased respiration is a hallmark of *Streptomyces* exploration. We observed significant upregulation of ATP synthesis and iron-sulfur cluster biogenesis in our transcriptional analyses here, and we have previously demonstrated that a functional copy of the alternative cytochrome *bd* oxidase is necessary to drive *S. venezuelae* exploration when grown next to yeast on YPD (14). A key biochemical property that distinguishes the cytochrome *bd* oxidase from the primary *aa*_3_ oxidase is a higher affinity for molecular oxygen (28–30). If respiration in exploring cultures is proceeding at a faster rate than classical vegetative growth, we may expect that local microenvironments of the colony could quickly become hypoxic, necessitating the need for stronger oxygen capture strategies. This may also present a self-imposed environment conducive to the function of the oxygen-sensitive Wbl-proteins. The intricate surface wrinkling patterns that we observe exclusively in exploring colonies may help cells respond to oxygen depletion by maximizing cell surface area for gas exchange. Indeed, increased colony wrinkling in response to oxygen depletion or impaired respiration appears to be a common response in microbial biofilm communities including those of *Pseudomonas aeruginosa* (31–33), *Bacillus subtilis* (34), *Candida albicans* (35), and *Aspergillus fumigatus* (36).

An unexpected outcome of probing the transcriptional program of explorer cells was the fascinating response of these colonies to glycerol supplementation. In the presence of glycerol, rates of exploratory growth readily outpaced traditional YP-grown explorers and colonies developed more complex wrinkling morphologies. Curiously, this enhancement appeared to be specific to glycerol; none of the other tested alternative carbon sources had the same exploration-promoting effect. This suggested that the role of glycerol during this growth mode may extend beyond simply acting as additional carbon substrate for biomass and energy production.

The architecture of exploring *Streptomyces* colonies is reminiscent of many bacteria biofilms. Notably, glycerol has been implicated in inducing or enhancing the establishment of bacterial biofilms in multiple systems. In the model organism *B. subtilis*, cultures supplemented with a combination of glycerol and manganese exhibit robust biofilm formation, with the histidine kinase KinD (for which no obvious homologue exists in *S. venezuelae*) being responsible for sensing and transducing this environmental signal (37). In *Listeria monocytogenes*, glycerol specifically induces biofilm formation at the air-liquid interface of aerobic broth cultures (38). Similar effects are seen in the pathogen *P. aeruginosa*, where growth on glycerol supports increased biofilm formation, in part through a resulting overproduction of Pel polysaccharide (39). In the yeast *C. albicans*, glycerol biosynthesis genes are upregulated during biofilm growth relative to planktonically-growing cells. Disruption of glycerol synthesis in this fungus leads to broad transcriptional changes, including a reduced capacity for biofilm formation and adherence, highlighting the role of this metabolite in the genetic regulation of biofilm development (40). Similarly, in the soil bacterium *Janthinobacterium lividum*, glycerol induces two phenotypic responses – increased biofilm formation (as evidenced through increased extracellular polysaccharide production) and overproduction of the purple pigment violacein (41). Production of an unknown – but highly conserved – pigment was also observed for the majority of glycerol-grown exploring *Streptomyces* cultures.

Given the strong response of exploring cultures to glycerol, it is worth considering how this observation is potentially reflective of the natural environment. The soil houses a particularly rich community of diverse microbes and is prone to large fluctuations in nutrient availability. In response to elevated glucose concentrations, yeasts like *S. cerevisiae* can produce and secrete glycerol as a by-product of fermentation (42, 43). This parallels what we know about exploration in *S. venezuelae*, where coculture with *S. cerevisiae* on YPD and the subsequent glucose depletion from the growth medium allows for a robust exploration response. It is conceivable that the accelerated rate of exploration in response to exogenous glycerol supplementation may be recapitulating an ecologically relevant strategy when encountering yeast under nutrient-rich growth conditions.

In all, we have shown that exploration requires a significant respiratory investment, and employs regulatory cascades that have some overlap but are largely distinct from those of classical development. How glycerol changes the exploration process, and what metabolic changes accompany exploration under all conditions, are questions of interest for future investigation.

## MATERIALS AND METHODS

### Strains, plasmids, media, and culture conditions

Strains, primers, and plasmids used in this study are listed in **Table S3, S4** and **S5** (provided in a supplementary data file). *S. venezuelae* NRRL B-65442 was grown in liquid MYM (1% malt extract, 0.4% yeast extract, 0.4% maltose) for overnight cultivation and solid MYM (2% agar) for spore stock generation and vegetative (non-exploratory) growth controls. Spore stocks for wild isolates were similarly prepared. For exploration experiments, 10 μL of an overnight culture of *S. venezuelae* (MYM, 10 mL) were spotted to solid YP medium (1% yeast extract, 2% peptone, 2% agar) additionally supplemented with 2% carbon source (e.g. dextrose, glycerol) and/or additional nutrients (sodium nitrate, sodium bisulfite, sodium sulfate; 10 μM, 100 μM, or 1 mM) where appropriate. In exploration experiments where the growth of multiple strains was being compared, overnight cultures were diluted with MYM to a normalized OD_600_. For experiments assessing the growth of wild *Streptomyces* isolates, due to frequent non-dispersed growth of different strains in liquid MYM, 5 μL of spore stock solutions were directly spotted to relevant solid media. All *Streptomyces* cultures were grown at 30 °C.

### *Construction of* S. venezuelae *mutants*

Gene deletions were generated using ReDirect technology (44). Coding sequences on a cosmid vector carrying large fragments (30-40 kb) of *S. venezuelae* genomic DNA were replaced by an *oriT*-containing apramycin [Δ*gylRFKD* (equivalent to Δ*vnz_06115-06130*); Δ*vnz_36165* (catalase-encoding gene); Δ*eshA* (Δ*vnz_35035*); Δ*sigQ* (Δ*vnz_22610*)] or hygromycin (Δ*bla* or Δ*gylRFKD*, for complementation control; see below) resistance cassette. In creating the *eshA* deletion strain, the coding sequence and flanking 2 kb up- and downstream sequences were PCR amplified and cloned into the pCR2.1-TOPO vector between the HindIII and SpeI restriction sites (as this region lacked an appropriate cosmid for deletion), before the *eshA* gene was targeted for replacement with the apramycin resistance cassette, as described above. Mutant cosmids/plasmid were introduced into the nonmethylating *E. coli* strain ET12567/pUZ8002 followed by conjugation into *S. venezuelae*. Resulting exconjugants were screened for double-crossover events and gene deletions were verified by PCR, using combinations of primers located upstream, downstream and internal to the deleted regions (see **Table S5**).

Complementation of the Δ*gylRFKD* mutation was accomplished by introducing a copy of the *S. venezuelae* cosmid Sv-3-E04 (containing a wild type copy of these genes plus associated upstream and downstream sequences) into the mutant strain. To facilitate conjugation and selection for cosmid integration into the *S. venezuelae* genome, the ampicillin resistance gene (*bla*) on the vector backbone was replaced with an *oriT*-containing hygromycin resistance cassette. To control for the additional up- and downstream sequences included on the cosmid, an analogous construct was generated by replacing the wild type *gylRFKD* coding region with an *oriT*-containing hygromycin resistance cassette. Both constructs were introduced into the *gylRFKD* mutant strain for phenotypic comparison with the wild type and mutant strains.

To generate the catalase overexpression construct, the coding sequence of *vnz_36165* was amplified from *S. venezuelae* genomic DNA with primers incorporating EcoRI and BamHI restriction enzyme recognition sites, which were subsequently used to clone the fragment directly downstream of the strong constitutive *ermE*\* promoter sequence in pMC500 (see Supplementary File 1). The promoter-gene fragment was then excised with KpnI and HindIII and was subcloned into the equivalently digested integrating plasmid pMS82 (see **Table S4**).

To generate the *sigQ* overexpression construct, a 487 bp fragment containing the *ermE*\* promoter was digested out from pIJ12251 using PvuII and EcoRV, and was subcloned into EcoRV-linearized pMS82. The coding sequence of *sigQ/vnz_22610* was subsequently amplified from the *S. venezuelae* genomic DNA with primers incorporating NdeI and XhoI restriction enzyme recognition sites (**Table S5**). The resulting amplicon was then cloned into the pMS82-*ermE*\*p construct following digestion by NdeI and XhoI (**Table S4**).

### Time-lapse videos of exploring colonies

Cells were inoculated onto exploration media, after which the plates were placed in a 30 °C incubator, on an Epson Perfection V800 Photo scanner that had been programmed to acquire one image every hour. The time-course images were then compiled to video format as sequential single frames.

### Solid media growth curves

Photos of growing colonies were taken at set time intervals. All image analyses were performed using ImageJ. Scale was established for each image by setting the diameter of the Petri dish to 10 cm. The perimeter of the colony was traced and the area of the captured region was determined.

### RNA isolation, library preparation, and cDNA sequencing

RNA was isolated as described previously (14), from two independent replicates of *S. venezuelae* grown on solid exploration medium for specified incubation times (2, 5, and 9 days of growth on YP). For all samples, ribosomal RNA (rRNA) was depleted using a Ribo-Zero rRNA depletion kit. cDNA and Illumina library preparation were performed using a NEBnext Ultra Directional Library Kit, followed by sequencing using unpaired-end 80 base-pair reads on the HiSeq Illumina platform. All bioinformatic analyses were carried out using packages available through the free open-access platform, Galaxy (usegalaxy.org). Reads were aligned to the *S. venezuelae* genome using Bowtie2 (45), then sorted, indexed, and converted to BAM format using SAMtools (46). Transcript level normalization and analyses of differential transcript levels were conducted using DESeq2 (47). RNA-seq data were submitted to the NCBI GEO repository and assigned the accession number GSE186259. COG analysis was performed using the open-source online tool EggNOG-mapper (48).

### Native PAGE assay for catalase enzyme activity

In-gel assays for catalase activity were conducted as described by Weydert and Cullen (49). Briefly, *S. venezuelae* cells were harvested from solid medium at timepoints and conditions of interest, before being resuspended in 1 mL of lysis buffer (100 mM NaCl, 5% glycerol, 10 mM Tris, pH 8; one cOmplete Mini EDTA-free protease inhibitor pellet (Roche) per 15 mL). Cell lysates were prepared by sonication (6 cycles of 15s of 40% duty followed by at least 15s on ice; Branson Sonifier Cell Disruptor 350) followed by centrifugation at >15,000 × *g* for 5 minutes at 4 °C. Protein concentration of lysate supernatants was determined by the Bradford assay (50). For each sample, 40 μg of total protein was loaded and resolved on an 8% non-denaturing acrylamide gel (19:1 acrylamide/bisacrylamide ratio) under constant voltage at 125 V for 2.5 hours. After separation, recovered gels were rinsed three times with Milli-Q water and then soaked in 0.003% hydrogen peroxide for 10 minutes with mild shaking. Following equilibration, gels were rinsed three times with Milli-Q water and then stained by the simultaneous addition of 10 mL of 2% ferric chloride and 10 mL of 2% potassium ferricyanide with gentle agitation. As soon as bands of clearing could be distinguished from the background staining of the gel, the excess stain solution was poured off and the gel was thoroughly washed with Milli-Q water to stop the reaction. As an additional loading control to compare protein concentrations between samples, 40 μg of protein was run under standard SDS-PAGE conditions on a 10% denaturing acrylamide gel (19:1 acrylamide/bisacrylamide ratio) under constant voltage at 125 V for 1 hour. Following separation, proteins were stained with Coomassie Brilliant Blue G-250.

## ACKNOWLEDGEMENTS

We would like to generously thank Gerry Wright for access to the Wright Actinomycete Collection, as well as Matt Hutchings and Thomas Mclean for helpful discussions. This work has been supported by a Natural Sciences and Engineering Research Council (NSERC) Discovery grant for MAE, a DFG Fellowship to TN (NE-2384/1-1), and an Ontario Graduate Scholarship and NSERC CGS-D doctoral scholarship to EMFS.

