## Supplementary material for "Exploratory growth in *Streptomyces venezuelae* involves a unique transcriptional program, enhanced oxidative stress response, and profound acceleration in response to glycerol": Tables S2, S3, S4, S5

**Table S2 – Effects of glycerol supplementation on exploring wild *Streptomyces* isolates**

| Strain | Accelerated growth rate in presence of glycerol[¥] | Induction of orange pigment secretion[¥] |
|---|---|---|
| WAC 666 | + | - |
| WAC 931 | - | + |
| WAC 1022 | + | - |
| WAC 1328 | + | + |
| WAC 1329 | + | + |
| WAC 2599 | + | - |
| WAC 3345 | + | + |
| WAC 4650 | + | Ambiguous* |
| WAC 4712 | + | + |
| WAC 4718 | + | + |
| WAC 4739 | + | + |
| WAC 5473 | - | + |
| WAC 5475 | - | + |
| WAC 5476 | - | + |
| WAC 5480 | - | + |
| WAC 5483 | + | + |
| WAC 5485 | + | + |
| WAC 5486 | - | + |
| WAC 5492 | - | + |
| WAC 5495 | - | + |
| WAC 6320 | + | + |

*In the presence of glycerol, WAC 4650 began producing a diffusible dark brown pigment (presumed to be melanin) that rendered assignment of orange pigmentation inconclusive.

[¥] The '+' indicates accelerated exploration and/or orange pigment production. The '-' indicates no enhanced exploration and/or no obvious orange pigmentation.

**Table S3 – Strains used in this study**

| Strains | Genotype/characteristics/use | Reference |
|---|---|---|
| ***Streptomyces*** | | |
| *S. venezuelae NRRL B-65442* | Wild type | (1) |
| WAC 666 | Wild *Streptomyces* isolate | Gift from G. Wright |
| WAC 931 | Wild *Streptomyces* isolate | Gift from G. Wright |
| WAC 1022 | Wild *Streptomyces* isolate | Gift from G. Wright |
| WAC 1328 | Wild *Streptomyces* isolate | Gift from G. Wright |
| WAC 1329 | Wild *Streptomyces* isolate | Gift from G. Wright |
| WAC 2599 | Wild *Streptomyces* isolate | Gift from G. Wright |
| WAC 3345 | Wild *Streptomyces* isolate | Gift from G. Wright |
| WAC 4650 | Wild *Streptomyces* isolate | Gift from G. Wright |
| WAC 4712 | Wild *Streptomyces* isolate | Gift from G. Wright |
| WAC 4718 | Wild *Streptomyces* isolate | Gift from G. Wright |
| WAC 4739 | Wild *Streptomyces* isolate | Gift from G. Wright |
| WAC 5473 | Wild *Streptomyces* isolate | Gift from G. Wright |
| WAC 5475 | Wild *Streptomyces* isolate | Gift from G. Wright |
| WAC 5476 | Wild *Streptomyces* isolate | Gift from G. Wright |
| WAC 5480 | Wild *Streptomyces* isolate | Gift from G. Wright |
| WAC 5483 | Wild *Streptomyces* isolate | Gift from G. Wright |
| WAC 5485 | Wild *Streptomyces* isolate | Gift from G. Wright |
| WAC 5486 | Wild *Streptomyces* isolate | Gift from G. Wright |
| WAC 5492 | Wild *Streptomyces* isolate | Gift from G. Wright |
| WAC 5495 | Wild *Streptomyces* isolate | Gift from G. Wright |

| | | |
|---|---|---|
| WAC 6320 | Wild *Streptomyces* isolate | Gift from G. Wright |
| E351 | *S. venezuelae vnz_06115-30::aac(3)IV* (glycerol uptake/metabolism operon) | This work |
| E352 | *S. venezuelae vnz_36165::aac(3)IV* (catalase-encoding gene) | This work |
| Δ*eshA* | *S. venezuelae vnz_35035::aac(3)IV* | This work |
| ***Escherichia coli*** | | |
| DH5α | Routine cloning | Invitrogen |
| SE DH5α | Highly-competent (Subcloning Efficiency™) DH5α cells | Invitrogen |
| BW25113/pIJ790 | Introducing mutations in cosmid DNA | (2) |
| ET12567/pUZ8002 | Generation of methylation-free plasmid DNA and conjugation into *Streptomyces* | (2) |

**Table S4 – Plasmids and cosmids used in this study**

| Cosmid/plasmid | Description | Reference |
|---|---|---|
| 2J14 | *S. venezuelae* cosmid carrying *vnz_36165* (*catC*) | Gift from M. Buttner |
| Sv-3-E04 | *S. venezuelae* cosmid carrying *vnz_06115-30* (glycerol uptake/metabolism operon) | Gift from M. Buttner |
| Sv-3-F11 | *S. venezuelae* cosmid carrying *vnz_22610* (*sigQ*) | Gift from M. Buttner |
| pIJ773 | Plasmid carrying the *aac(3)IV-oriT* cassette | (2) |
| pIJ10700 | Plasmid carrying the *hyg-oriT* cassette | (3) |
| pIJ10701 | Plasmid carrying the *bla-hyg-oriT-bla* cassette for recombination with cosmid backbone | (3) |
| pIJ12551 | Plasmid carrying the strong *Streptomyces* promoter *ermE**promoter (p) | (4) |
| pMC500 | Plasmid carrying the strong *Streptomyces* promoter *ermE**p | (5) |
| pMC300 | *S. venezuelae vnz_36165* (*catC*) coding sequence cloned downsteam of *ermE**p in pMC500 | This work |
| pMS82 | Integrative cloning vector: *hyg*, *oriT*, *int* ΦBT1, *attP* ΦBT1 | (6) |
| pMC301 | *ermE**p-*vnz_36165* (*catC*) sequence from pMC300 subcloned into pMS82 | This work |
| pMC302 | *ermE**p-*vnz_22610* (*sigQ*) cloned into pMS82 | This work |
| pCR2.1-TOPO | Cloning vector used for creating *eshA* deletion plasmid | Invitrogen |
| pCR2.1-TOPO_*vnz_35035* | pCR2.1-TOPO carrying *vnz_35035* (*eshA*)and 2 kb up- and downstream flanking regions | This work |
| pCR2.1-TOPO_Δ*vnz_35035* | Vector for deletion of *vnz_35035* (*eshA*), pCR2.1-TOPO_*vnz35035*::*aac(3)IV* | This work |

**Table S5 – Oligonucleotides used in this study**

| Name | Sequence (5' to 3')* | Use |
|---|---|---|
| *vnz_06115* ReD FWD | CTGTCATGTGTCATCCAAGGTCGGCATCGACCGA TAGTG***ATTCCGGGGATCCGTCGACC*** | Creation and confirmation of the Δ*vnz_06115-30* (glycerol uptake/metabolism operon) mutation |
| *vnz_06130* ReD REV | GGAGCCGCCCCTGAACCGTCCCTGAAGCCGGGG GAGTTA***TGTAGGCTGGAGCTGCTTC*** | Creation and confirmation of the Δ*vnz_06115-30* (glycerol uptake/metabolism operon) mutation |
| *vnz_06115* Up FWD | ATGAGCTCCCTGGGATGCGA | Confirmation of the Δ*vnz_06115-30* (glycerol uptake/metabolism operon) mutation |
| *vnz_06115* In REV | CGCTCGTTCAAGCGACTGGAT | Confirmation of the Δ*vnz_06115-30* (glycerol uptake/metabolism operon) mutation |
| *vnz_06130* Down REV | TCTGCGCGCGGAGTGTTG | Confirmation of the Δ*vnz_06115-30* (glycerol uptake/metabolism operon) mutation |
| *vnz_36165* ReD FWD | CTGAGCACGCGATCCGCCCCGTCCGGAAGGATT TCCATG***ATTCCGGGGATCCGTCGACC*** | Creation and confirmation of the Δ*vnz_36165* (*catC*) mutation |
| *vnz_36165* ReD REV | GTCGACCGGGCCCGGGACAGCGGGGGTTGAGC GGGATCA***TGTAGGCTGGAGCTGCTTC*** | Creation and confirmation of the Δ*vnz_36165* (*catC*) mutation |
| *vnz_36165* Up FWD | TCGTCGGCCGTGAGGTGCAC | Confirmation of the Δ*vnz_36165* (*catC*) mutation |
| *vnz_36165* In REV | GACTCCTGGGGTGCGGGGTC | Confirmation of the Δ*vnz_36165* (*catC*) mutation |
| *vnz_36165* OE FWD EcoRI | CATCATGAATTCCTAGCCAAGGCACACCACGG | Creation of *vnz_36165* (*catC*) overexpression construct |
| *vnz_36165* OE REV BamHI | CATCATGGATCCGGAGGCGGGTTGTGGATCGC | Creation of *vnz_36165* (*catC*) overexpression construct |
| *sigQ* ReD FWD | GGACCGAAACACACTCCAACCGACGGGGGGCG CCAGATG***ATTCCGGGGATCCGTCGACC*** | Creation and confirmation of the *vnz_22610* (*sigQ*) mutation |
| *sigQ* ReD REV | CCGTTTCCCCCGTGAGCCGCAGGGTCCCCCCGAC CTTCA***TGTAGGCTGGAGCTGCTTC*** | Creation and confirmation of the *vnz_22610* (*sigQ*) mutation |
| *sigQ* Up FWD | GACAGTCGGCGGACAACGC | Confirmation of the *vnz_22610* (*sigQ*) mutation |
| *sigQ* In REV | GTCGCTGATCCGCTCCCAGG | Confirmation of the *vnz_22610* (*sigQ*) mutation |
| *sigQ* OE FWD NdeI | GGTCGGCATATGACACACTCCAACCGACGGG | Creation of *vnz_22610* (*sigQ*) overexpression construct |
| *sigQ* OE REV XhoI | CATCATCTCGAGCAGGGTCCCCCCGACCTTC | Creation of *vnz_22610* (*sigQ*) overexpression construct |
| *eshA* flank FWD HindIII | CAGTGTAAGCTTGTTGTGCAGCTCCACGAC | Vector creation for generating the *vnz_35035* (*eshA*) mutation |
| *eshA* flank REV | CGTTGCACTAGTCGTACATGCTCGACTCGTTG | Vector creation for generating the |

| | | |
|---|---|---|
| SpeI | | *vnz_35035* (*eshA*) mutation |
| *eshA* ReD FWD | CCGGTCACCACCCCCCTGCGAAGGAGAGCCCGC CCCATG***ATTCCGGGGATCCGTCGACC*** | Creation and confirmation of the *vnz_35035* (*eshA*) mutation |
| *eshA* ReD REV | GACTGTCATGGCCGCCGTGGCGGCGGCCCGGG ACCTCTA***TGTAGGCTGGAGCTGCTTC*** | Creation and confirmation of the *vnz_35035* (*eshA*) mutation |
| *eshA* WT check | GCCGCTGCAGAAGCAGAAC | Confirmation of the *vnz_35035* (*eshA*) mutation |
| *eshA* KO check | CGCAGTAGCAGTCGTCGACG | Confirmation of the *vnz_35035* (*eshA*) mutation |
| blaF | ***CCCTGATAAATGCTTCAATAATATTGAAAAAGG AAGAGTA*** | Amplification of *bla-hyg-oriT-bla* from pIJ10701 for cosmid backbone replacement |
| blaR | ***AATCAATCTAAAGTATATATGAGTAAACTTGGT CTGACAG*** | Amplification of *bla-hyg-oriT-bla* from pIJ10701 for cosmid backbone replacement |

*Cassette-specific sequences are bolded and italicized. Restriction enzyme recognition sequences are underlined.


1. Gomez-Escribano JP, Holmes NA, Schlimpert S, Bibb MJ, Chandra G, Wilkinson B, Buttner MJ, Bibb MJ. 2021. *Streptomyces venezuelae* NRRL B-65442: genome sequence of a model strain used to study morphological differentiation in filamentous actinobacteria . J Ind Microbiol Biotechnol.
